# Identification and comparative analysis of long non-coding RNAs in the brain of fire ant queens in two different reproductive states

**DOI:** 10.1101/2021.08.18.456372

**Authors:** Cheng-Hung Tsai, Tzu-Chieh Lin, Yi-Hsien Chang, Huai-Kuang Tsai, Jia-Hsin Huang

**Author notes:** **Corresponding authors** Correspondence to Jia-Hsin Huang or Huai-Kuang Tsai.

## Abstract

**Background:** Many long non-coding RNAs (lncRNAs) have been extensively identified in many higher eukaryotic species. The function of lncRNAs has been reported to play important roles in diverse biological processes, including developmental regulation and behavioral plasticity. However, there are no reports of systematic characterization of long non-coding RNAs in the fire ant *Solenopsis invicta*.

**Results:** In this study, we performed a genome-wide analysis of lncRNAs in the brains of *S. invicta* from RNA-seq. In total, 1,393 novel lncRNA transcripts were identified in the fire ant. In contrast to the annotated lncRNA transcripts having at least two exons, novel lncRNAs are monoexonic transcripts with a shorter length. Besides, the transcriptome from virgin alate and dealate mated queens were analyzed and compared. The results showed 295 differentially expressed mRNA genes (DEGs) and 65 differentially expressed lncRNA genes (DELs) between virgin and mated queens, of which 17 lncRNAs were highly expressed in the virgin alates and 47 lncRNAs were highly expressed in the mated dealates. By identifying the DEL:DEG pairs with high association in their expression (Spearman’s |*rho*| > 0.8 and *p*-value < 0.01), many DELs were co-regulated with DEGs after mating. Furthermore, several remarkable lncRNAs (*MSTRG*.*6523, MSTRG*.*588*, and *nc909*) that were found to associate with particular coding genes may play important roles in the regulation of brain gene expression in reproductive transition in fire ants.

**Conclusion:** This study provides the first genome-wide identification of *S. invicta* lncRNAs in the brains in different reproductive states and will contribute to a fuller understanding of the transcriptional regulation underpinning reproductive changes.

## Background

The red imported fire ant (*Solenopsis invicta* Buren) is one of the notorious invasive species globally [1]. The introduction of *S. invicta* has been reported in several countries, including the USA, New Zealand, Taiwan, and Japan. The fire ant invasion has caused not only substantial economic losses but also a drastic impact on the agricultural systems and ecological environments.

*S. invicta* is a eusocial insect with large colonies containing interdependent divisions of individuals of all different developmental stages and multiple castes [2]. This ant species could form a polygynous colony, in which multiple queens live together, within the subterranean nests. In a mature polygynous nest, dealate mated queens and virgin alate queens perform distinct reproductive statuses associated with different physiology and behaviors [3]. Recently, Calkins et al. [4] reported that the differential expression of some protein-coding genes between virgin and mated queens are linked to nutritional and immune processes occurring in queens’ behavioral transitions. However, this previous study focused on protein-coding genes while many regulated transcripts including long non-coding RNAs (lncRNAs) were uncharacterized yet. Long noncoding RNAs (lncRNAs) refer to a group of noncoding RNAs that are arbitrarily characterized as a transcript of more than 200 nucleotides in length and lack coding potential in the eukaryotic cells [5]. LncRNAs possess distinguishing features, including shorter length, fewer exon numbers, and lower expression levels compared to protein-coding genes [6, 7]. In recent years, many lncRNAs have been identified in Hymenoptera insects [8–10]. It is noteworthy that some lncRNAs were found to play a potential regulatory relationship with protein-coding genes in the adult caste transition between worker and gamergates in *Harpegnathos saltator* [10] and in the behavioral transition from nurses to foragers of *Apis mellifera* [9]. So far, there are no reports about the identification of lncRNAs in the fire ant *S. invicta*.

Here, we report the utilization of RNA-seq data to identify and characterize lncRNAs in the brains of *S. invicta*. In this study, we identified a high-confident set of 1,393 novel lncRNA transcripts and further characterized the basic features of lncRNA transcripts in the fire ant brain. In conjugation with annotated lncRNAs, a total of 65 lncRNAs were significantly differentially expressed in the fire ant brains during the virgin to mated status transition. Collectively, genome-wide annotation and characterization of *S. invicta* brain lncRNAs pave the way for further studies to identify genetic and molecular mechanisms of phenotypic and behavioral plasticity in the fire ants.

## Results

### Identification of novel lncRNAs in fire ant brains

The *S. invicta* genome is not yet completely annotated based on the updated chromosome assembly. However, larger scaffolds were recently assembled and obtained more predicted genes, including 1,211 lncRNAs, than the previous assembly [11]. In order to comprehensively understand the relationship between lncRNA and dealation in *S. invicta*, we developed a pipeline to identify putative, unannotated lncRNAs and further examine the expression differences of all lncRNAs between virgin alate and mated dealate. The flowchart of processing steps in our pipeline is shown in Figure 1.

**Figure 1.**
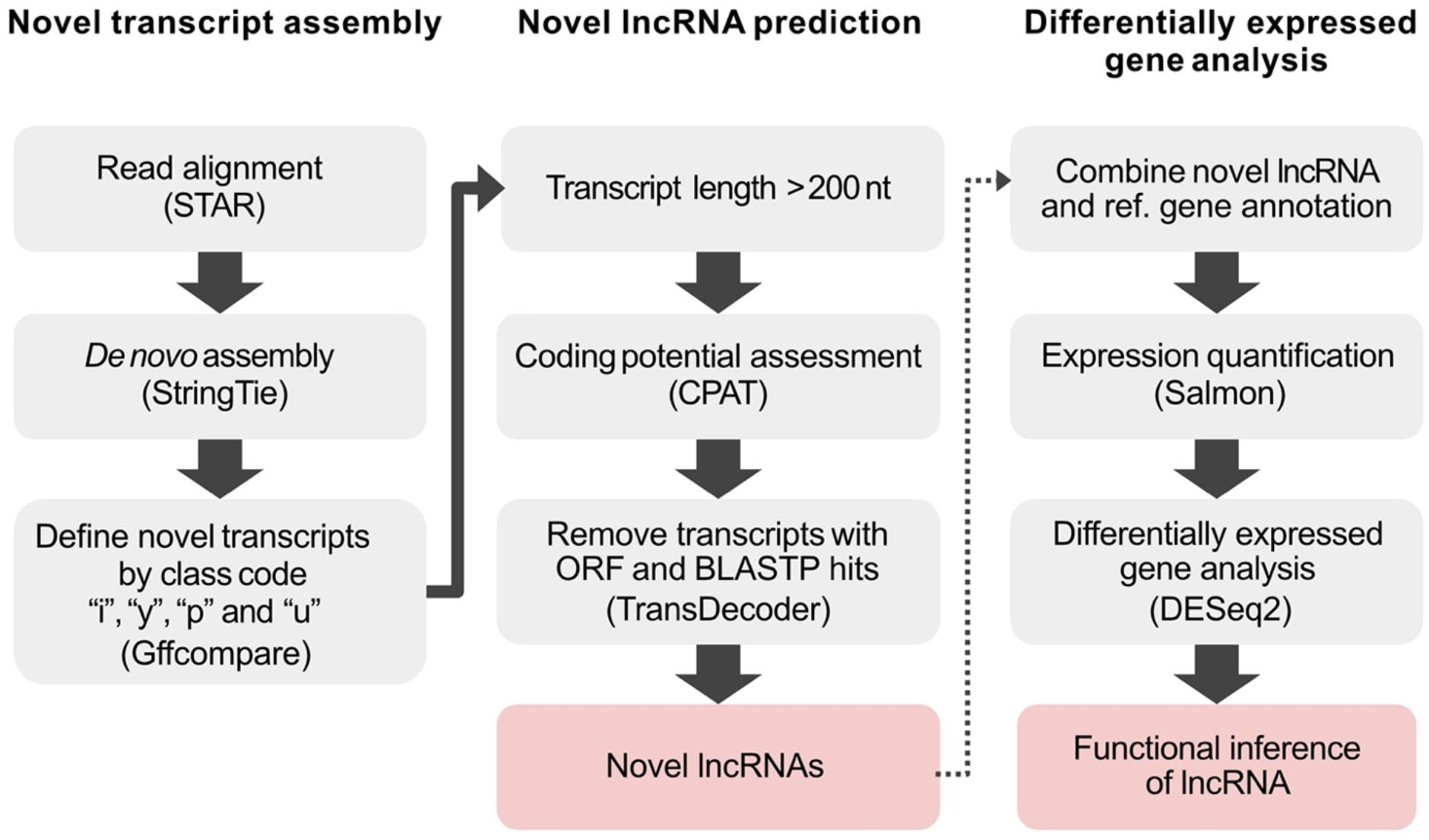
Flowchart of the pipeline used to identify novel lncRNAs and differential gene analysis from the RNA-seq in *S. invicta*.

The first part of our pipeline was to identify novel transcripts from RNA-seq data. We used a total of 32 RNA-seq libraries collected from virgin alate and mated dealate queen brains of *S. invicta*. Approximately 100 million clean reads were obtained after removing the contaminated and low-quality reads; the mapping rate of RNA-seq libraries ranged from 97.78% to 99.22% by STAR [12]. The alignment files were then followed by a *de novo* assembly using Stringtie [13]. Filtering through the class codes (i.e., i, y, p, and u) defined by Gffcompare [14], we further identified 1,829 novel transcripts.

Next, multiple filtering steps were performed to identify putative lncRNAs from our novel transcripts collection. First, the novel transcripts having over 200 nt in length were selected for coding potential or open-reading frame (ORF) evaluations. Second, we employed the Coding Potential Assessment Tool (CPAT) [15] to estimate the coding probability according to the transcript sequence features. Two ant species, *Camponotus floridanus* and *Harpegnathos saltator*, having well-curated lncRNA genes, were used to determine the optimal coding probability cutoff as 0.224 for identifying non-coding transcripts (Figure 2A). Then, TransDecoder [16] determines potential coding regions by identifying the ORF from transcripts. Candidate transcripts were then fed to homology search via BLASTP. The transcripts which were denoted as candidate coding sequences by TransDecoder and BLASTP were then discarded. Using this pipeline, we finally identified a set of 1,393 novel lncRNA transcripts derived from 1,393 loci in the fire ant genome (See Additional file 1).

**Figure 2.**
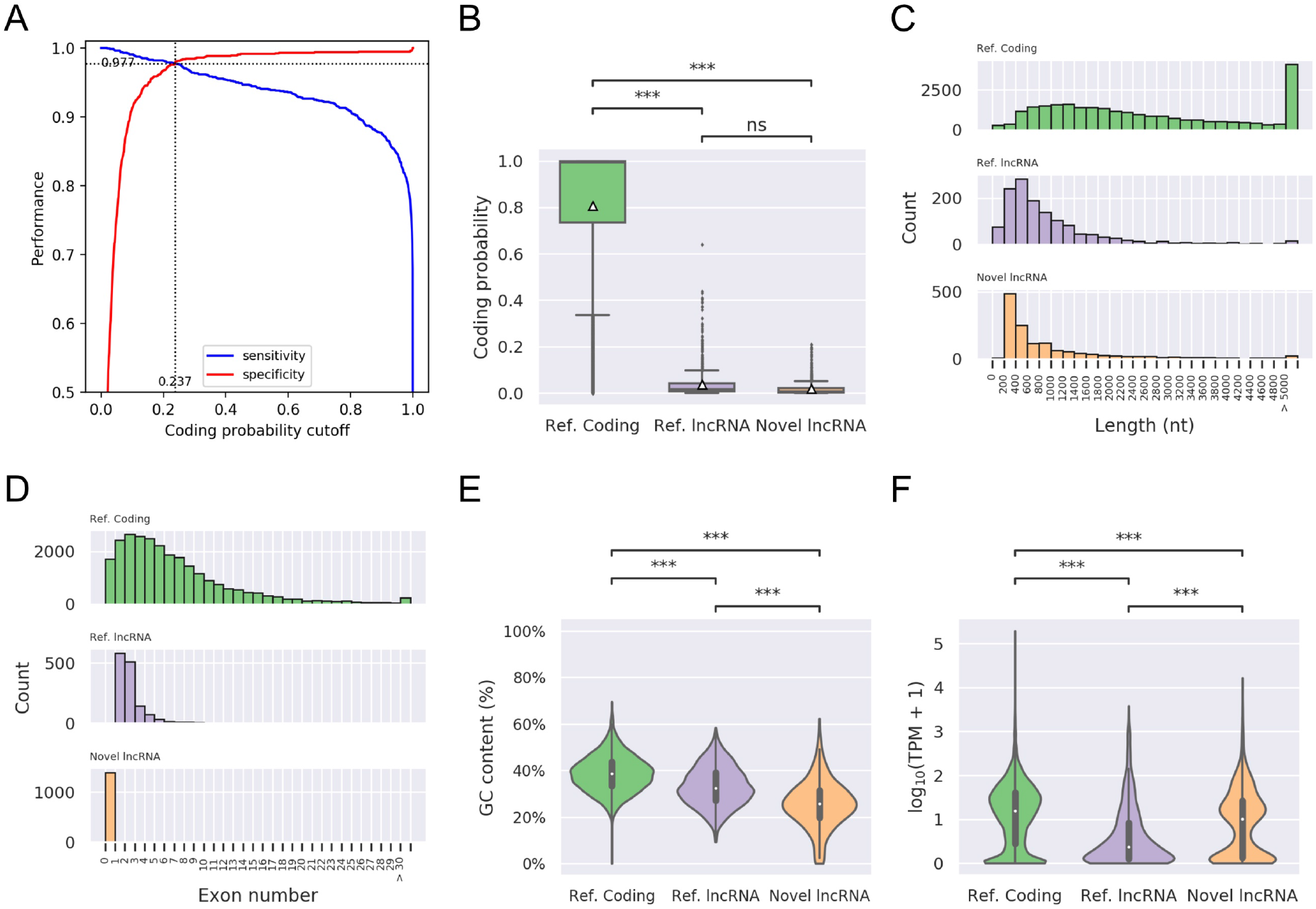
Sequence features and expression analysis of different transcript types. (A) Two ROC curves for determination of CPAT threshold of coding probability using annotated lncRNAs and mRNAs of *S. invicta*. The accuracy of model performance was 0.977 as the coding probability at 0.237. (B) Coding probability distribution of known protein coding and lncRNA, and novel lncRNA transcripts. (C) Length distribution of known protein coding and lncRNA, and novel lncRNA transcripts. (D) Distribution of the number of exons in the coding and lncRNA transcripts. (E) Distribution of the GC content in the coding and lncRNA transcripts. (F) Expression level distribution of known coding and lncRNA, and novel lncRNA transcripts. Expression level is indicated by log_10_(TPM+1). The *p-*values were calculated using Mann-Whitney test and following post hoc HSD test, *** *p*-value < 0.0001.

We further examined the general characteristics of our newly identified lncRNA transcripts. First, we compared the coding potential among reference coding genes, reference lncRNAs and novel lncRNAs. As expected, novel lncRNAs are significantly lower in coding potential than reference coding genes, but no differences in contrast to reference lncRNAs. The average coding probability of coding gene, reference lncRNAs and novel lncRNA transcripts are 0.086, 0.036 and 0.018, respectively (Figure 2B). For the length of transcripts, novel lncRNAs show a similar distribution compared to reference lncRNAs and are relatively shorter than coding genes (Figure 2C). The average length of novel lncRNAs is 1,025 nucleotides while the longest lncRNA transcript contains 8,436 nucleotides. Next, we examined the number of exons contained in each lncRNA transcript. Note that all of the reference lncRNAs are spliced and constitute at least two exons (Figure 2D). In contrast, all of our newly identified lncRNA transcripts have a single exon. In addition, GC content of fire ant lncRNAs is significantly lower than that of coding genes while newly identified lncRNAs have lowest GC content (Figure 2E). Of the lncRNA characteristics, lower GC content in lncRNAs is linked to some biological functions [17].

Based on the expression quantification using salmon [18], we profiled the protein coding genes, reference lncRNAs, and novel lncRNAs predicted by this study for virgin alate and mated dealate queens in Figure 2F. Overall, the expression levels (TPM) of the protein coding genes were significantly higher than those of lncRNAs in the fire ant brain. However, the overall expression of novel lncRNAs were significantly higher than that of reference lncRNAs. About half of the newly identified lncRNAs were highly-expressed (TPM ≥ 10) in the fire ant queen brain. These results suggested that some of lncRNAs would play important roles in the brain function of *S. invicta*.

### Gene expression analysis and DE genes identification

To investigate the transcriptional changes between virgin alate to dealate mated queen brains, we identified the differentially expressed (DE) genes based on the criteria with log_2_(fold change) ≥ 1 and FDR < 0.01 by DESeq2 [19] (see Additional file 2). Of those protein coding genes (mRNAs), 155 were found to be upregulated and 140 were found to be downregulated during the transition from virgin alate to dealate mated states (Fig. 3A). Next, we conducted a hierarchical clustering analysis using average TPM values of these 395 DEGs, which classified the eight queen brain samples into two groups: virgin alate and mated deleted, as expected (Fig. 3B).

**Figure 3.**
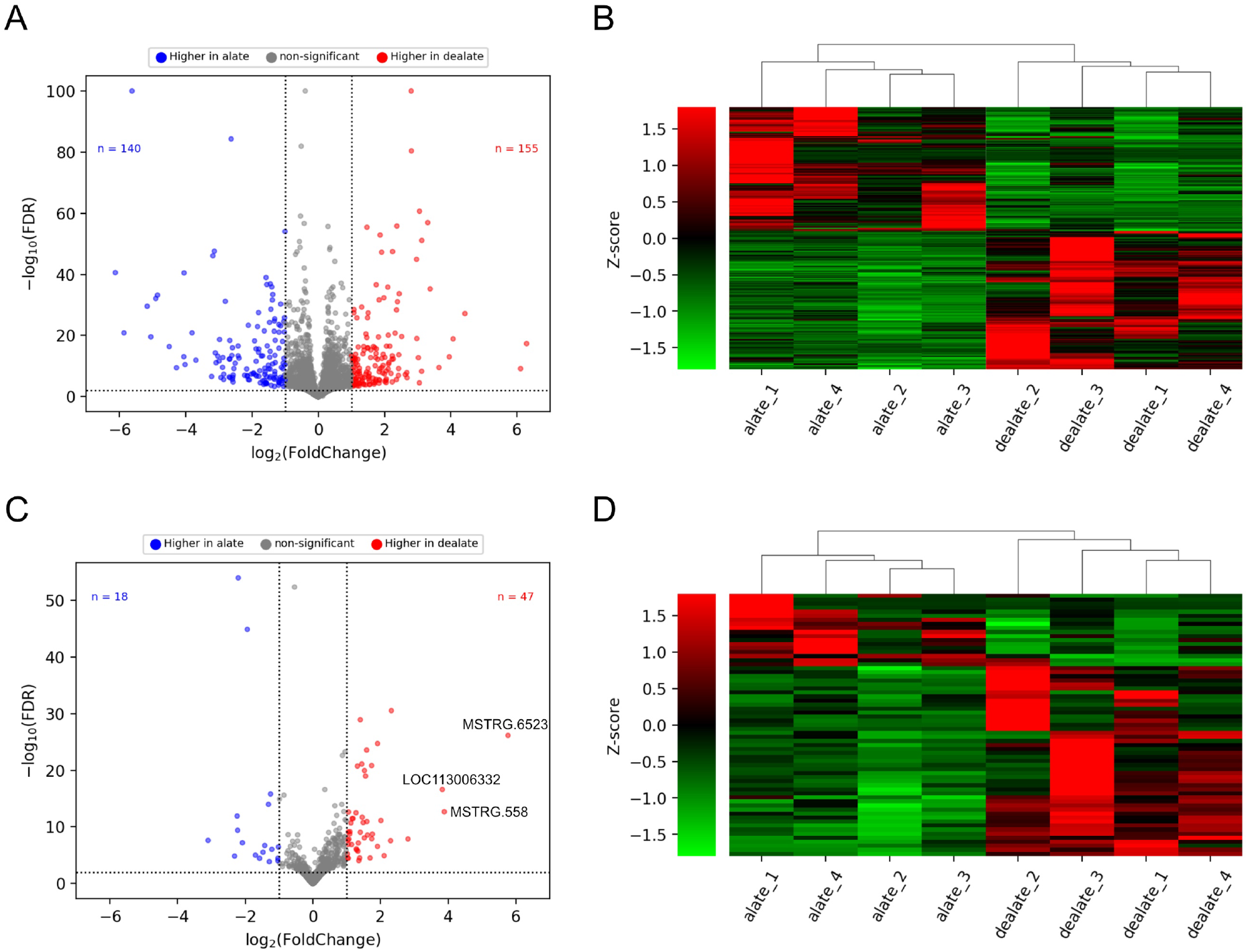
Coding gene (mRNA) expression profiles of virgin alate and mated dealate queen brains. (A) Volcano plot analysis of coding gene expression variation. Blue and red points correspond to the significantly enriched mRNA (DEGs) in the virgin and mated queens respectively with the |log_2_(fold-changes)| ≥ 1 and FDR < 0.01. (B) Hierarchical clustering of the 295 DEGs in the 8 independent biological replicates. For each biological replicate, the values were calculated by averaging four technical runs of RNA-seq libraries. Red represents the high expression levels and Green represents the low expression levels. (C) Volcano plot analysis of lncRNA expression variation. Blue and red points correspond to the significantly enriched lncRNA (DELs) in the virgin and mated queens respectively with the |log_2_(fold-changes)| ≥ 1 and FDR < 0.01. (D) Hierarchical clustering of the 65 DELs in the 8 independent biological replicates. For each biological replicate, the values were calculated by averaging four technical runs of RNA-seq libraries. Red represents the high expression levels and Green represents the low expression levels.

Of the DE lncRNAs (DELs), a total of 65 lncRNAs were considered to be differentially expressed between the virgin alate and dealate mated queen brains (log_2_FC ≥ 1 and FDR < 0.01) by DESeq2. In total, 28 lncRNAs were annotated in the reference genome while 37 lncRNAs were newly identified in the current study. Among these DELs, 18 were found upregulated in the virgin alate group while 47 were found upregulated in the dealate mated group (Fig. 3C). Furthermore, top three most upregulated lncRNAs showed larger increase (at least 16-fold) in the dealate mated queen brains in contrast to those in the virgin alate queen brains. Hierarchical clustering of virgin alate and dealate mated queen brain samples based on 65 DELs revealed a clear separation of the two reproductive states (Fig. 3D).

### Co-regulation of DE lncRNAs and mRNAs

In the previous study, the adult reproductive transition between virgin and mated queen brains is accompanied by major changes in particular protein-coding gene expression with transcriptome verification and validation [4]. In order to explore the potential function of these DELs, we employed the correlation analysis of expression profiles between DEGs and DELs in the virgin alate and dealate mated queen brain samples, respectively (Fig. 4A). The common principle in such correlation analysis is that two transcripts display consistently correlated across many samples (here are 16 samples in virgin and mated groups respectively) of their expression levels, and thereby, are determined to be co-regulated. Such co-regulation may suggest that either the two transcripts are regulated by common factors or one may regulate another’s expression. According to the most notable theme of lncRNA function appears to act as the *cis* or *trans* regulation on the mRNA gene expression in the eukaryotic cells [20, 21], we exploited the highly correlated DEL:DEG pairs to infer the potential regulation of genes during reproductive status transition.

**Figure 4.**
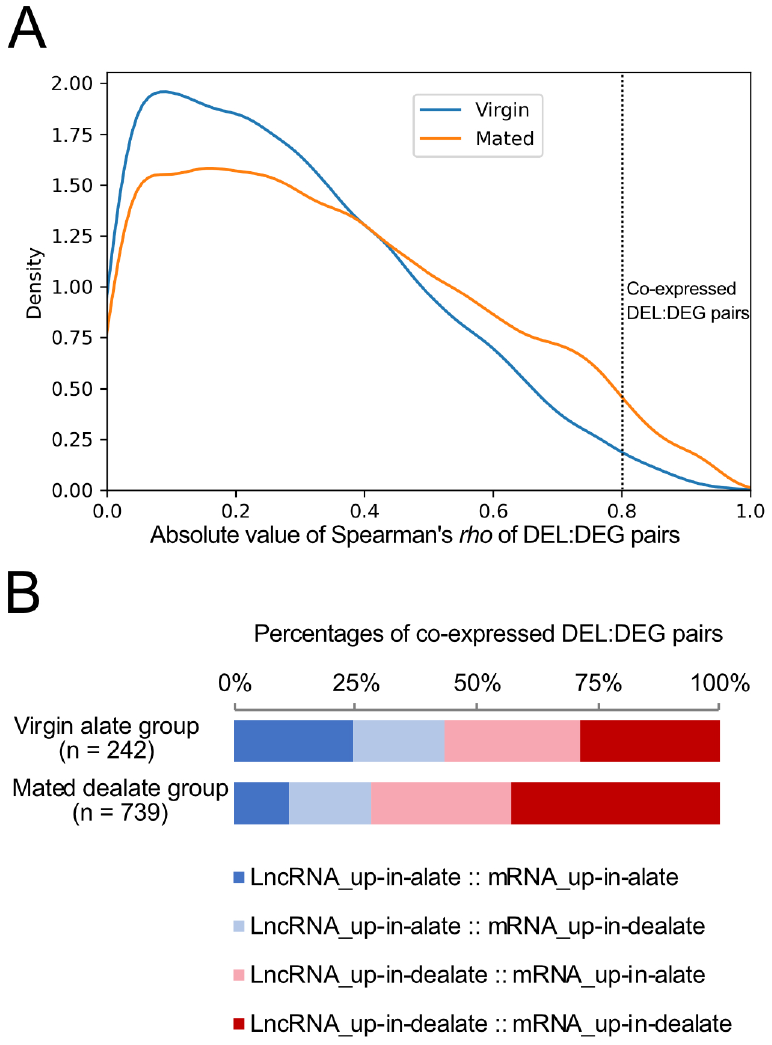
Correlation of expression of lncRNAs and protein-coding genes. (A) Density plot of correlations of expression for differentially expressed lncRNAs (DELs) and differentially expressed mRNAs (DEGs) in the virgin alate and mated dealate samples respectively. The co-regulated DEL:DEG pairs have a highly correlated profile of expression (Spearman’s *rho* > 0.8, *p*-value < 0.01). (B) Proportions of different pairs of the co-regulated DEL:DEG in the virgin and mated groups.

We calculated the Spearman correlation coefficient between DELs and DEGs, and obtained the genes with a Spearman’s |*rho*| > 0.8 and *p*-value < 0.01 as the co-regulated mRNAs of lncRNAs. In the virgin group, a total of 242 DEL:DEG pairs were highly correlated (Fig. 4B). While in the mated group, a total of 739 DEL:DEG pairs were highly correlated. In the following sections, we delved into the functional significance of the several highly enriched lncRNAs in the mated dealate queen brains identified in this study.

### Functional assessment of differentially expressed lncRNAs

First, we examined the top one DEL, MSTRG.6523, which is a new annotated lncRNA in this study, and found that 7 DEGs were highly associated (Fig. 5A). In particular, the protein coding gene *LOC105192919* (encoded *hexamerin 1*) caught our attention because its expression was the only mRNA showed strongly negative correlation with MSTRG.6523 in the mated dealated samples (Spearman’s *rho* = -0.818, *p*-value = 1.083×10^−8^, Fig. 5B). One of the notable mechanisms of lncRNA molecules in the transcriptional regulation is through RNA-DNA interactions, and thereby inhibiting the expression of target genes [22]. Consequently, we applied conducted the Triplextor program [23] to conduct the RNA:DNA:DNA triplex prediction by using MSTRG.6523 and the genomic sequences nearby the *LOC105192919* locus. Notably, a cluster of triplex target sites (TTSs) of *MSTRG*.*6523* lies within the first intron region of *hexamerin 1* (scaffold NW_020521759.1: 281973-282016, Fig. 5C). It is noteworthy that this region of triplex clusters occurred at a high frequency of 15 positions with GA-rich DNA sequences (Fig. 5D).

**Figure 5.**
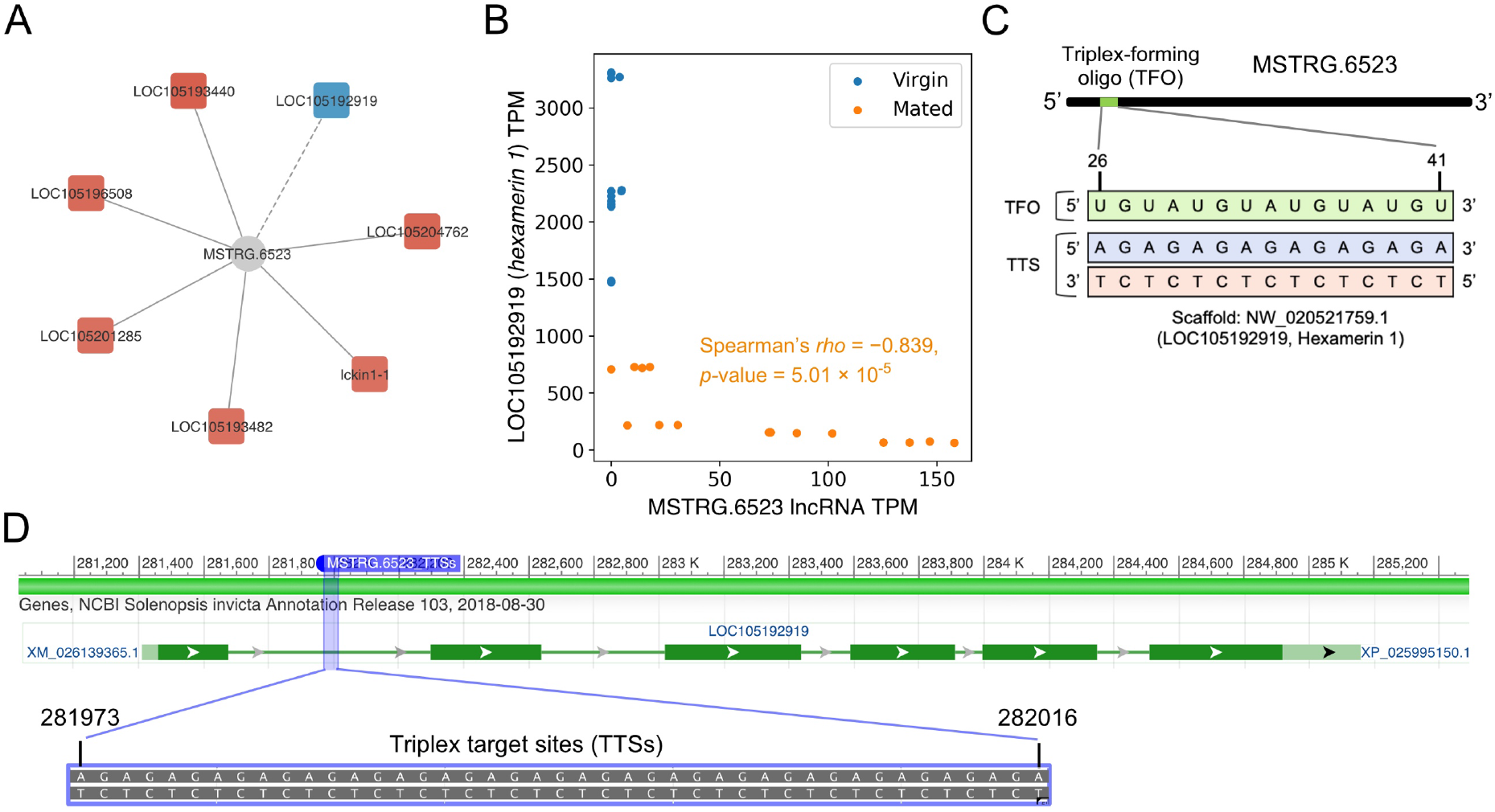
Relationships between *MSTRG*.*6523* lncRNA and protein-coding gene expression. (A) The co-regulation network of *MSTRG*.*6523* lncRNA and other DEGs in the mated dealate samples. Solid lines and red color in DEGs denote the positive correlation and dash lines and blue color in DEGs denote negative correlation. (B) The expression levels of *MSTRG*.*6523* lncRNA (x axis) and the protein-coding gene LOC105192919 encoded as *hexamerin 1* (y axis) correlate in mated dealate queen brains. Each dot represents one biological sample (virgin, n = 16; mated, n = 16). Correlation coefficient (*rho*) and *p*-value from Spearman correlation of mated samples is indicated. (C) Triplex-forming oligonucleotide (TFO) motif within *MSTRG*.*6523* lncRNA (green) identified using Triplexator software to target triplex sites (TTSs) in *hexamerin 1* locus (blue and red) to form a RNA:DNA:DNA triplex. (D) Predicted triplex binding sites of *hexamarin 1* locus (purple box) with *MSTRG*.*6523* lncRNA and the AG-rich sequences are presented in the first intron of *hexamarin 1* gene.

Next, we examined the top 2 of DELs, *MSTRG*.*558*, which is up-regulated in the mated dealate and found that its expression was strongly correlated with the expression of two chymotrypsin-inhibitor (CI) like genes, *LOC105200276* and *LOC105200273* (Fig. 6A and 6B). After looking for other co-exregulated DEL:DEG pairs, we noted that these two CI genes were significantly correlated with several DELs (Fig. 6C). Interestingly, the up-regulated DELs in the mated dealate were positively correlated with these two CI genes while the down-regulated DELs were all negatively correlated. For example, a down-regulated DELs, *LOC1052639*, in the mated dealate showed significantly negative correlations with both CI-like genes, LOC105200276 and LOC105200273 (Fig. 6D and 6E). Such reversed relationships suggest a good co-regulated relationship between lncRNAs and CI genes during the mating transition.

**Figure 6.**
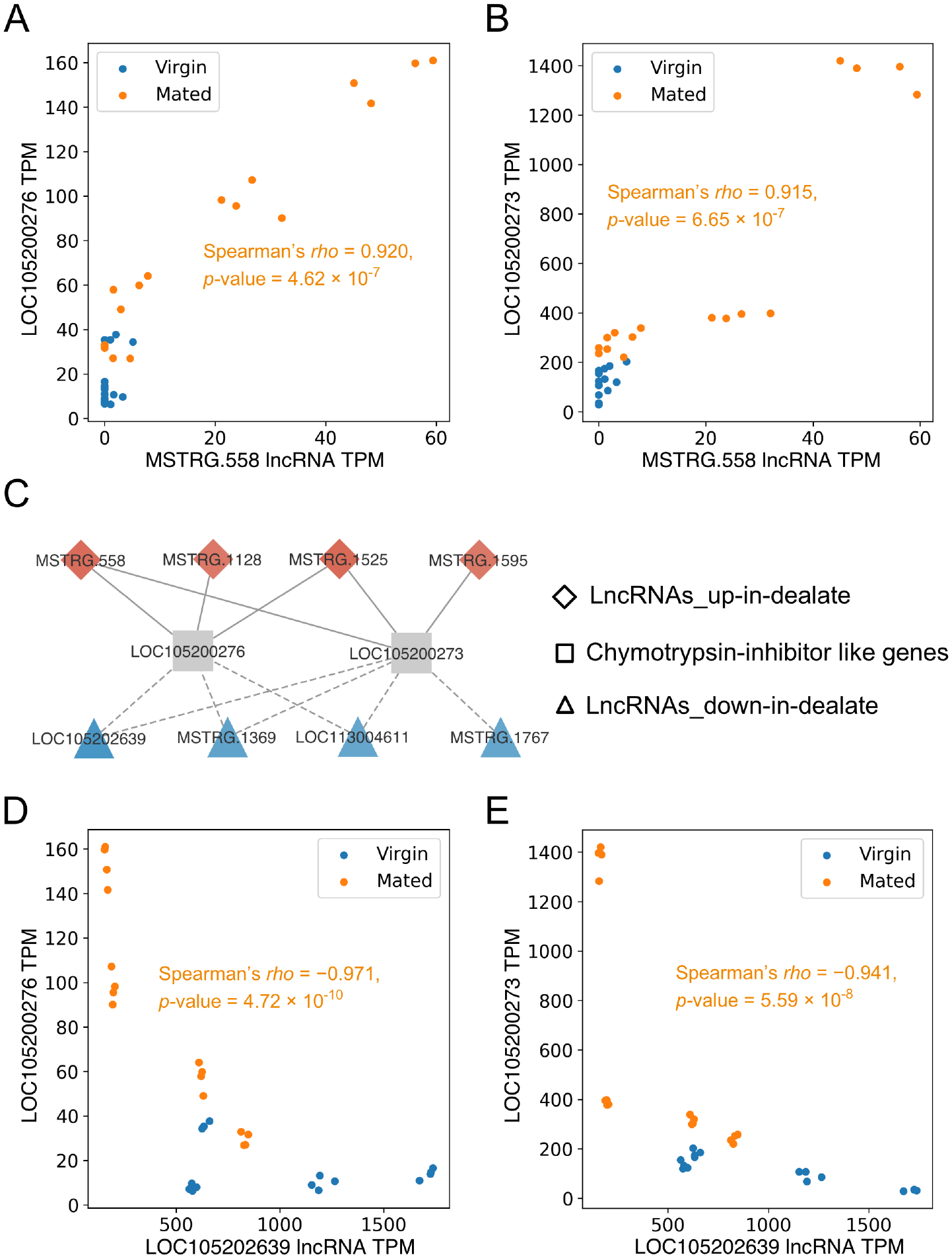
Co-regulatory module between lncRNAs and chymotrypsin-inhibitor (CI) genes. Scatter plots of *MSTRG*.*558* lncRNA expression across all fire ant queen brains with two enriched-in-mated-dealate CI genes, *LOC105200276* (A) and *LOC105200273* (B). Correlation coefficient (*rho*) and *p*-value from Spearman correlation of mated samples is indicated respectively. (C) A co-regulation network of DELs and two CI genes. Solid lines and red color in DEGs denote the positive correlation and dash lines and blue color in DEGs denote negative correlation. Plotting the expression of *LOC105200276* (D) and *LOC105200273* (E) against that of *LOC105202639* lncRNA across all virgin and mated samples showed negative correlations significantly.

In addition, a significant elevation of *LOC105199067* lncRNA (designated *nc909* in the previous study) expression in the mated dealate queens has been validated independently from field-collected samples by quantitative real-time PCR [4]. Here, we also examined the probable functions of *nc909* lncRNA by analyzing its co-regulated DEGs. It is noteworthy that the co-regulated networks of nc909 and associated DEGs showed distinct patterns between virgin alate and mated dealate groups (Fig. 7A). Also, the co-regulated DEGs were almost not overlapped between two reproductive states, except for *LOC105200275*. A highly positive correlation in the expression profiles between *nc909* and *LOC105200275*, which is encoded as a CI gene, across both alate and dealate samples was observed (Spearman’s *rho* = 0.946, *p*-value = 2.88 × 10^−16^, Fig. 7B). This implies that expression of *nc909* is coordinately regulated with DEGs during the reproductive transition from virgin to mated states. As a large amount of *nc909* transcripts was existed in the mated dealates compared to virgin alate queen brains, it caught our attention to further investigate its regulatory role in the mated dealate queen brains. Particularly in the mated dealates, the expression of *nc909* transcripts had negative correlations with most of the co-regulated DEGs, either the down-regulated-in-mated DEGs such as *LOC1052019089* (Fig. 7C) or the up-redulated-in-mated DEGs such as *LOC106195192* and *LOC105197201* (Fig. 7D and 7E, respectively). With the notable mechanism of lncRNA function to suppress gene expression via recruitment of chromatin modulators such as polycomb repressive complex 2 (PRC2) [22, 24], we hypothesized that *nc909* acts as an attenuator to control the gene expression in the queen brains after mating.

**Figure 7.**
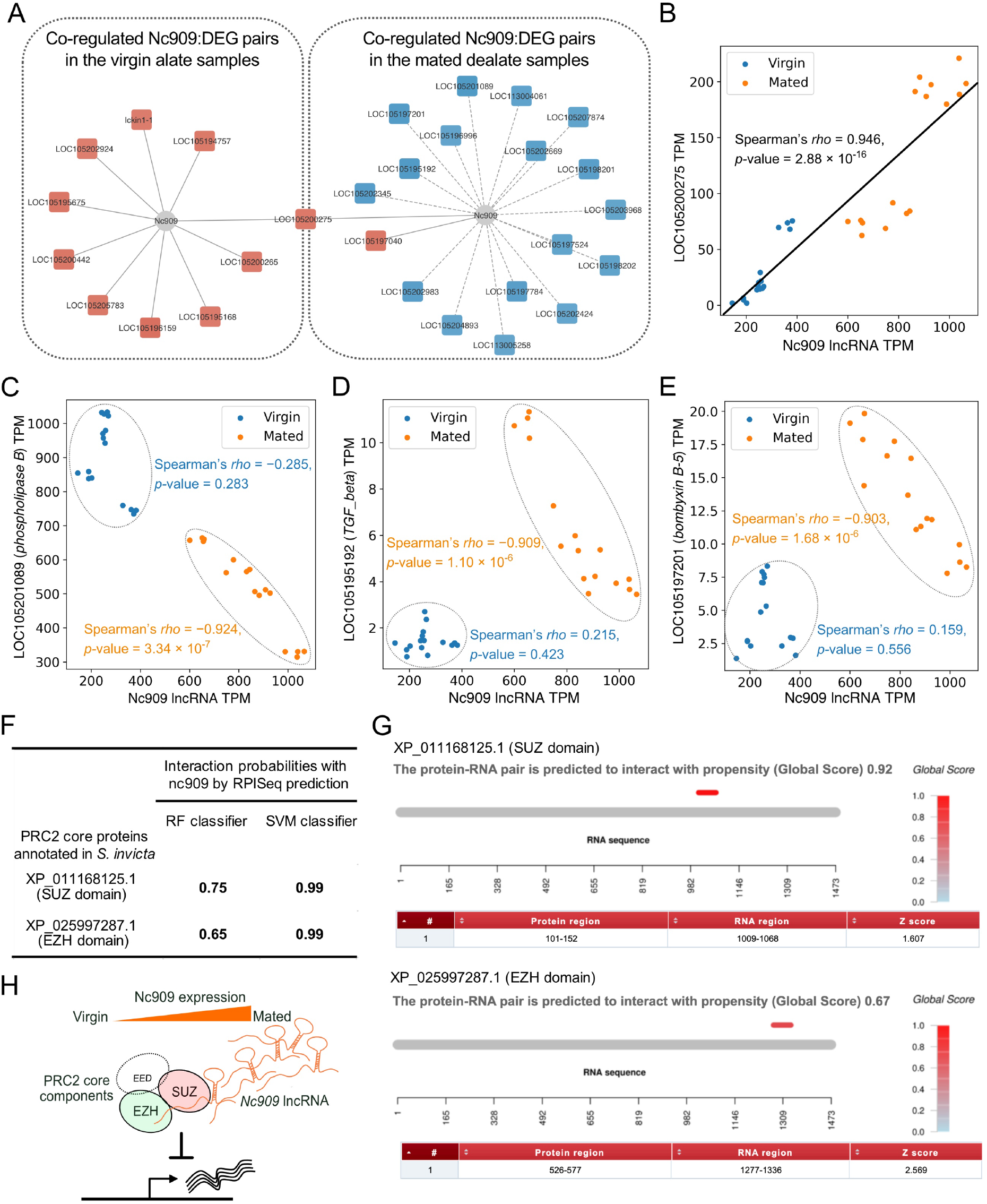
Nc909 lncRNA expression associated with DEGs. (A) A co-regulation network of *nc909* lncRNA with DEGs in the virgin and mated groups, respectively. Solid lines and red color in DEGs denote the positive correlation and dash lines and blue color in DEGs denote negative correlation. (B) The expression profile of *nc909* and *LOC105200275* (chymotrypsin inhibitor gene) shows a significantly positive correlation across both virgin and mated samples. *Nc909* lncRNA expression has significantly negative correlations with *LOC105201089* (C), *LOC105195192* (D), and *LOC105197201* (E) across the mated samples (orange) but no correlation across virgin samples (blue). (F) RPISeq predictions on the *nc909* lncRNA and two annotated proteins of PRC2 core components in *S. invicta*. (G) The catRAPID outputs for the signal localization of *nc909* lncRNA and two PRC2 core proteins binding propensity prediction with Global Score value, signal localization plot which denotes the localization of the signal along the RNA sequence colored according to the Global Score value, and a table with the information of the binding loci with a Z-score representing the interaction propensity in contrast to benchmark value of a pool of RNA-binding proteins. (H) Proposed model of the up-regulated *nc909* lncRNA transcripts interacting with PRC2 to inhibit gene expression in the mated dealate queens brains.

In order to test this hypothesis, we used the computational approaches to examine the likelihood of RNA-protein interaction using RNA-Protein Interaction Prediction (RPISeq) web service [25] for *nc909* lncRNA and the core proteins of PRC2 in *S. invicta*. We have identified two annotated *S. invicta* gene products, XP_011168125.1 with SUZ domain and XP_025997287.1 with EZH domain, which were reported to be the core proteins of PRC2 [26]. The interaction probabilities predicted by RPISeq for *nc909* lncRNA and two PRC2 core proteins were summarized in Fig. 7F. Since the RPISeq prediction with probabilities greater than 0.5 were considered positive interacting [25], nc909 is most likely to interact with these two *S. invicta* PRC2 proteins based on their high probability scores from both classifiers. Besides, we employed the catRAPID, an algorithm to predict the potential RNA-protein interacting sites [27, 28] from catRAPID interactions with larger RNAs, the interaction of nc909/PRC2 interactions are further supported with the discriminative power of 0.92 and 0.67 were identified to interact with XP_011168125.1 and XP_025997287.1, respectively (Fig. 7G).

## Discussion

In this paper, publicly available RNAs-eq data from fire ant queen brains were collected and re-analyzed to excavate brain related transcripts. We have built a pipeline to *de novo* assemble and predict 1,829 novel transcripts, specifically for non-coding lncRNAs. Of note, all of the reference lncRNAs are annotated with at least two exons (Figure 2D). In fact, single-exon lncRNAs have been identified in almost all the eukaryotic genomes. All of our newly identified lncRNA transcripts having a single exon indeed complement the previous studies in the annotation of lncRNAs in *S. invicta* genome. Besides, pseudoalignment methods such as salmon [18] have been reported to show a better performance in lncRNA quantification with full transcriptome annotation [19]. Thus, we followed the recommended strategy and quantified the gene expression of each RNA-seq run by salmon, instead of using the expression profiles from StringTie in our lncRNA discovery pipeline. Since the expression level of novel lncRNAs were significantly higher than reference lncRNAs (Fig. 2F), we assumed that newly lncRNAs play a major role in the regulation of the fire ant brain.

Previously, the authors had identified only 19 DE coding genes (DEGs) between virgin alate and mated dealate queen brains [4]. By contrast, we identified 295 DEGs in total (Fig. 3A) and 16 out of the19 DEGs were found in our results. Nevertheless, the comparison of expression changes in the 16 DEGs confirms a consistent trend between previous and our methods (Pearson’s *r* = 0.965, *p*-value < 0.0001 in terms of fold-change similarity). The reason for having different numbers of DEGs identified in our results could be accounted for by the newer version of genome assembly (GCF_000188075.2) and gene annotation file as well as up-to-date transcript quantification tools used for the RNA-seq analysis in this study.

In the co-expression analysis between DEGs and DELs (Fig. 4), larger amounts of highly correlated DEL:DEG pairs in the mated dealate samples could be partly due to the larger number of DELs in the mating condition than those in the virgin condition. However, the highest proportion of DEL:DEG pairs in the mated group was the differentially expressed lncRNAs and mRNAs in the mated dealate (Fig. 4B). This is an interesting observation, which indicates the lncRNAs and mRNAs were co-regulated during the transition of mating states and worth investigating further.

A key contribution of this study is to provide functional inferences of several important lncRNAs in the fire ant queen brains. Nowadays increasing evidences point to the presence of lncRNAs acting as a transcriptional regulator in the nucleus by different mechanisms, including interaction with chromatins and RNA-binding proteins [24, 29]. Experiments on the chromatin-binding maps of lncRNAs have revealed that GA-rich sequences are the preferred binding motifs to help these RNAs to target the chromatin [25, 30]. A recent study has demonstrated that the high frequency of triplex loci in the human *WWOX* gene is important for *PARTICLE* lncRNA binding to regulate its expression [31]. It is noteworthy that expression of *hexamerin 1* was significantly reduced in the mated dealate queen brains of *S. invicta* with experimentally validations [4, 32]. We therefore inferred that *MSTRG*.*6523* lncRNA is upregulated in the queen brains during the reproductive transition after mating and potentially binds to the *hexamerin 1* intron to suppress its expression (Fig. 5). The probable mechanism of *MSTRG*.*6523* lncRNA interacting with *hexamerin 1* intron DNA via RNA:DNA:DNA triplex formation is worthy to be taken into account for future study.

In addition, a fluctuation of *nc909* lncRNA expression in the fire ant queen brains was experimentally validated to link to the mating and social context in the previous study [4], but its function remained unexplored. Here we applied *in silico* analyses to provide novel insights into *nc909* lncRNA functions. In contrast to virgin alate queen brains, the expression level of *nc909* lncRNA was increased drastically in the dealate mated queen brains (Fig. 7B). Surprisingly, a negative correlation between *nc909* lncRNA and most co-expressed DEGs could be only observed during the elevation of *nc909* lncRNA expression levels (Fig. 7A). This scenario suggests that a potential mechanism of *nc909* lncRNA to regulate gene expression may be due to its high abundance. One well-known human lncRNA MALAT1 (metastasis-associated lung adenocarcinoma transcript 1) is highly abundant in the nucleus and suppresses targeting genes via interacting with the PRC2 complex [33]. Following this idea, we here proposed that a plausible role of *nc909* lncRNA serves as a negative regulator in gene expression by interacting with the transcriptional repressors such as PRC2 core proteins (Fig. 7F) in the mated dealate queen brains (Fig. 7H). Further experiments are needed to be performed to explore the specific regulatory mechanisms of *nc909* lncRNAs in the queen brains of *S. invicta*.

Besides, the CI genes are tightly regulated to inactivate the serine proteases to avoid the hazard of excessive peptidase activities in the living cells. Correspondingly, we found a coding gene *LOC105200925* as the most enriched chymotrypsin 2 like proteases with 64-fold increased expression in the mated dealates without any of correlated lncRNAs. However, our correlation study suggests a fine regulation between CI genes and several lncRNAs during mating transition (Fig. 6). Taken together, we suspect that lncRNAs and CI genes are co-regulated to maintain cellular homeostasis since several protease genes were also up-regulated in the mated dealate brains.

The major limitation of current study on the inference of lncRNA function is the lack of high quality assembly of the *S. invicta* genome when analyzing the RNA-seq data. Although we employed analysis using the updated version of *S. invicta* genome (Si_gnH version GCF_000188075.2) than that in the original paper, the number of scaffolds in the current assembly version remains 66,904. In fact, most genes including annotated genes and novel lncRNAs are located in the short scaffolds, and thereby, cause difficulties to infer the putative function of those lncRNAs using the physical genomic distance to study their *cis*-regulatory function. In addition, many of protein-coding genes, which were highly associated with DEL in their expression levels, were annotated as uncharacterized proteins. Despite the challenges remaining in the functional study of lncRNAs in the fire ant, we conducted *in silico* prediction for potential roles of lncRNAs with investigation of the co-regulated lncRNA:mRNA networks thoroughly.

## Conclusion

In this report, we have presented the first genome-wide identification of novel lncRNAs in the fire ant S. *invicta*. Firstly, we identified 1,393 novel lncRNAs in the *S. invicta* with a custom bioinformatics pipeline. In addition, 18 and 47 lncRNAs were significantly enriched in virgin alate and mated dealate queen brains, respectively. Lastly, we elucidated several remarkable lncRNAs (*MSTRG*.*6523, MSTRG*.*588*, and *nc909*) and their roles to associate with specific coding genes, which play important biological functions in the mated queen brains. To sum up, this study provides novel insights of lncRNAs for further studies to obtain a deeper understanding of *S. invicta* brain function during reproductive maturation.

## Methods

### RNA-seq data acquisition

The raw reads of a total 32 RNA-seq dataset were downloaded from NCBI Sequence Reads Archive (SRA) with accession number SRP126736 [4]. The RNA-seq datasets include fire ant alate virgin and dealate mated queen transcriptomes. All libraries were prepared with single-end protocol and on average contains 4 million reads.

### RNA-seq processing and lncRNA identification

High-quality clean reads were obtained by clipping adapters and low-quality reads with Trimmomatic [34] version 0.38. Each library was individually mapped to the *S. invicta* genome (NCBI Si_gnH version GCF_000188075.2) using STAR [12] version 2.7 with the default parameters. The resulting alignment files were used as input for transcript assembly using StingTie [13] version 2.1.1 with *S. invicta* reference annotation of NCBI Release 103. All Stringtie output files were merged into a single unified transcriptome assembly using Stringtie merge option. Gffcompare [14] was employed to compare the resulting unified transcriptome assembly (GTF format) to the reference annotation, and transcripts with class code *i, y, p, u* were obtained for lncRNA prediction analysis. To identify the lncRNA, the assembled transcripts with a minimum length of 200 nt were subjected to the coding potential assessment tool (CPAT) [15] to determine their coding probability. We used the lncRNAs identified from other two ant species *Camponotus floridanus* and *Harpegnathos saltator* [10] for benchmark tests on CPAT and set the cutoff threshold as 0.224. Then, we discarded the remaining transcripts with any open-reading frame and potential homologs to other species predicted by TransDecoder [16] and BLASTP to annotate novel lncRNA transcripts.

### Gene expression and differentially expressed genes

We used Salmon [18] version 1.2.0 to quantify the gene expression with default parameters. Read count for each transcript was subjected for differential analysis using DESeq2 [19] in R environment. Any gene with zero read counts in less than 3 samples of mated or virgin groups was filtered out for comparison. The differentially expressed genes (DEGs) were identified at least two fold differences with 0.01 false discovery rate (FDR). The TPM of all replicates in each sample were averaged, followed by a z-transform to normalize the expression differences among all DEGs. Hierarchical clustering analysis of the averaged TPM was done in a python environment by seaborn API.

### LncRNA-mRNA Correlation test

The TPM values of all genes from Salmon were used to analyze their association relationship between lncRNA and coding genes using Spearman’s correlation test. The cutoff of Spearman’s correlation coefficient |*rho*| > 0.8 and *p*-values < 0.01 was applied to define co-regulated DEL:DEG pairs.

### Prediction of lncRNA:DNA triplexes

Triplexator [23] is a computational framework for the *in silico* prediction of triplex structures by forming Hoogsteen base-pairing rules [35]. To predict lncRNA:DNA triplex sites, we employed the Triplexator program with parameters to constrain the maximal error less than 20%, minimal guanine content of triplex target sites as 20%, at most 3 constitutive errors, and limited the length of TTS-TFO pair as 15.

## Abbreviations

lncRNA: long non-coding RNA
DEG: differentially expressed mRNA gene
DEL: differentially expressed lncRNA gene
ORF: open-reading frame
CPAT: Coding Potential Assessment Tool
TPM: Transcripts per million
FDR: False discovery rate
FC: Fold change
CI: Chymotrypsin-inhibitor
RPISeq: RNA-Protein Interaction Prediction
PRC2: Polycomb repressive complex 2
EZH: Enhancer of zeste homolog
SRA: Sequence Reads Archive
TFO: Triplex-forming oligo
TTS: Triplex target site
nt: nucleotide

## Declarations

### Authors’ contributions

CHT conceived the study, implemented the bioinformatics pipelines and prepared the initial draft of the manuscript. TCL curated the data and carried out the bioinformatics analysis. YHC assisted the data curation and conducted preliminary experiments. HKT contributed to the experimental design and edited the manuscript. JHH conceived and designed the research, carried out the bioinformatics analysis, interpreted the results, prepared the initial draft and edited the manuscript. All authors read and approved the final manuscript.

## Acknowledgements

Not applicable.

## Availability of data and materials

The raw reads of a total 32 RNA-seq dataset were downloaded from NCBI Sequence Reads Archive (SRA) with accession number SRP126736 (https://www.ncbi.nlm.nih.gov/sra?term=SRP126736).

## Ethics approval and consent to participate

Not applicable.

## Consent for publication

Not applicable.

